# Enhanced Piezoelectric Performance of PVDF-TrFE Nanofibers through Annealing for Tissue Engineering Applications

**DOI:** 10.1101/2024.08.16.608345

**Authors:** Maksym Krutko, Holly M. Poling, Andrew E. Bryan, Manju Sharma, Akaljot Singh, Hasan A. Reza, Kathryn A. Wikenheiser-Brokamp, Takanori Takebe, Michael A. Helmrath, Greg M. Harris, Leyla Esfandiari

## Abstract

This study investigates bioelectric stimulation’s role in tissue regeneration by enhancing the piezoelectric properties of tissue-engineered grafts using annealed poly(vinylidene fluoride-trifluoroethylene) (PVDF-TrFE) scaffolds. Annealing at temperatures of 80°C, 100°C, 120°C, and 140°C was assessed for its impact on material properties and physiological utility. Analytical techniques such as Differential Scanning Calorimetry (DSC), Fourier-Transform Infrared Spectroscopy (FTIR), and X-ray Diffraction (XRD) revealed increased crystallinity with higher annealing temperatures, peaking in β-phase content and crystallinity at 140°C. Scanning Electron Microscopy (SEM) showed that 140°C annealed scaffolds had enhanced lamellar structures, increased porosity, and maximum piezoelectric response. Mechanical tests indicated that 140°C annealing improved elastic modulus, tensile strength, and substrate stiffness, aligning these properties with physiological soft tissues. In vitro assessments in Schwann cells demonstrated favorable responses, with increased cell proliferation, contraction, and extracellular matrix attachment. Additionally, genes linked to extracellular matrix production, vascularization, and calcium signaling were upregulated. The foreign body response in C57BL/6 mice, evaluated through Hematoxylin and Eosin (H&E) and Picrosirius Red staining, showed no differences between scaffold groups, supporting the potential for future functional evaluation of the annealed group in tissue repair.

## 1. Introduction

Bioelectricity has shown to be a regulator of cell and tissue properties and influence a broad spectrum of biological processes such as embryogenesis, tissue repair, and wound healing [1–6]. Bioelectric stimuli can influence favorable changes in cellular phenotype enhancing regulation of ECM deposition and upregulating genes involved in tissue repair in both excitable and non-excitable cell types such as neurons and Schwann cells (SCs) respectively [7–9]. Post tissue damage, a bioelectric flux serves as a primary repair signal guiding the regenerative cascade of subsequent inflammation, cell migration, new blood vessel formation, and tissue remodeling [10–12]. Thus, it has been theorized that incorporating electrical stimuli into the next generation of scaffolds for tissue repair can improve regenerative outcomes. Conductive and piezoelectric biomaterials are at the forefront of this innovation, offering improved integration and functionality compared to traditional materials [2, 13, 14]. Conductive scaffolds can be utilized to electrically stimulate tissue for regeneration; however, their reliance on an external power source to apply electric fields can pose challenges such as complications with wire interfaces, risks of overheating, and potential electrical hazards [15–18]. In contrast, piezoelectric biomaterials are more versatile and can harness their ability to generate electrical energy under mechanical stress and vice versa [19]. Notably, piezoelectric scaffolds can be activated through bodily movements in numerous tissue types including cellular traction forces and can also synergize with non-invasive activation techniques such as ultrasound or shockwaves [20–27]. The versatility of piezoelectric scaffolds makes them effective in tissue repair strategies in various physiological systems.

Electrospinning, a fabrication technique where polymers are electrically and mechanically poled to create tunable nanofiber scaffolds, advantageously enhances the piezoelectric nature of piezopolymers [28–32]. Leveraging electrospinning technology, we have previously shown aligned PVDF-TrFE nanofiber scaffolds are suitable for peripheral nerve repair and skin innervation [28–32]. Despite these advantages, electrospun scaffolds face challenges with mechanical strength and low piezoelectric sensitivity, making them less effective in stimulating non-excitable tissue nor able to handle high load-bearing tissues such as bone [33–35]. Although piezoelectric composites, that join piezoelectric ceramic and polymer constituents, address these concerns, their clinical translation for soft tissue applications is hindered by potential metallic ion toxicity and corrosiveness [36]. Thermal post processing techniques such as annealing may remedy these constraints without further poling or doping, provide favorable outcomes through modifying molecular microstructure to enhance piezoelectric coefficients (d33) and potentially enhance material properties [37–40]. This trend has also been observed with annealed electrospun polycaprolactone (PCL) and piezoelectric Poly l-Lactic Acid (PLLA) scaffolds, which exhibited enhanced crystallinity, mechanical strength, and chemical stability [41–43]. Additionally, in another study, polyvinylidene fluoride (PVDF) and PVDF-TrFE films were annealed between 100 °C and 140 °C, increasing piezoelectricity of polymers while promoting favorable morphologies for cellular attachment [44–46]. However, Mao et al. has recently shown that the most suitable annealing temperature of PVDF-TrFE to achieve high piezoelectricity varies from case-to-case basis dependent on fabrication of piezoelectric material of interest, annealing temperature and time [47].

This study thus, systematically explores the effects of annealing treatments on electrospun nanofiber PVDF-TrFE scaffolds in the context of morphological, mechanical, and piezoelectric properties. We specifically examine relevant changes in bulk Young’s modulus, substrate stiffness, crystallinity, and other emergent material properties due to annealing treatments. We further assess the biocompatibility of these annealed scaffolds, showcasing enhanced cellular responses through metabolic assays with murine fibroblast and Schwann Cell (SC) lines. Additionally, we evaluated protein production related to cell-extracellular matrix interactions using western blot analysis and investigated gene expression through RNA-sequencing to understand the impact on cell attachment and repair phenotype of SCs on annealed scaffolds. Our findings demonstrated that the annealed PVDF-TrFE nanofibers have the potential to be utilized as a biomimetic material with the enhanced porosity, mechanical strength, substrate stiffness, and piezoelectric properties for tissue repair. We also unveil their ability to contribute to the enhanced proliferation and repair phenotypes of SCs within the same framework. Additionally, our proof of concept analysis of subcutaneous implantations of annealed and untreated or control scaffolds in a mouse model demonstrated no difference in foreign body responses highlighting the potential for annealed PVDF-TrFE scaffolds in future functional implementation and testing.

## 2. Results

### 2.1 Thermal analysis

Thermogravimetric analysis (TGA) illustrates the weight loss of PVDF-TrFE scaffolds (Figure 1A), depicting the thermal stability of the material. Beyond 406°C, decomposition occurs, and complete degradation of electrospun PVDF-TrFE happens at approximately 445°C. Differential scanning calorimetry (DSC) provides important information on a polymer’s thermal properties (Figure 1B). Notably, two prominent endothermic peaks are observed at 99°C and 154°C in non-treated (NT) or control PVDF-TrFE scaffolds, corresponding to the curie temperature and melting temperature, respectively [48–51]. Scaffolds annealed at 80°C, 100°C, 120°C, and 140°C revealed similar crystallization ranges which were used to determine total crystallinity fraction percentage (%) (Eq 1). Where in the expression for overall crystallinity, enthalpy of melting or Δ*H*_*m*_ is obtained by integrating the area under the melting peak of each DSC curve located at around 144°C, and Δ*H_g_* is the melting enthalpy for fully crystalline PVDF-TrFE (70/30) (26.1 J/g) depicted in Eq 1 [52].

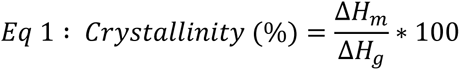

**Figure 1:**
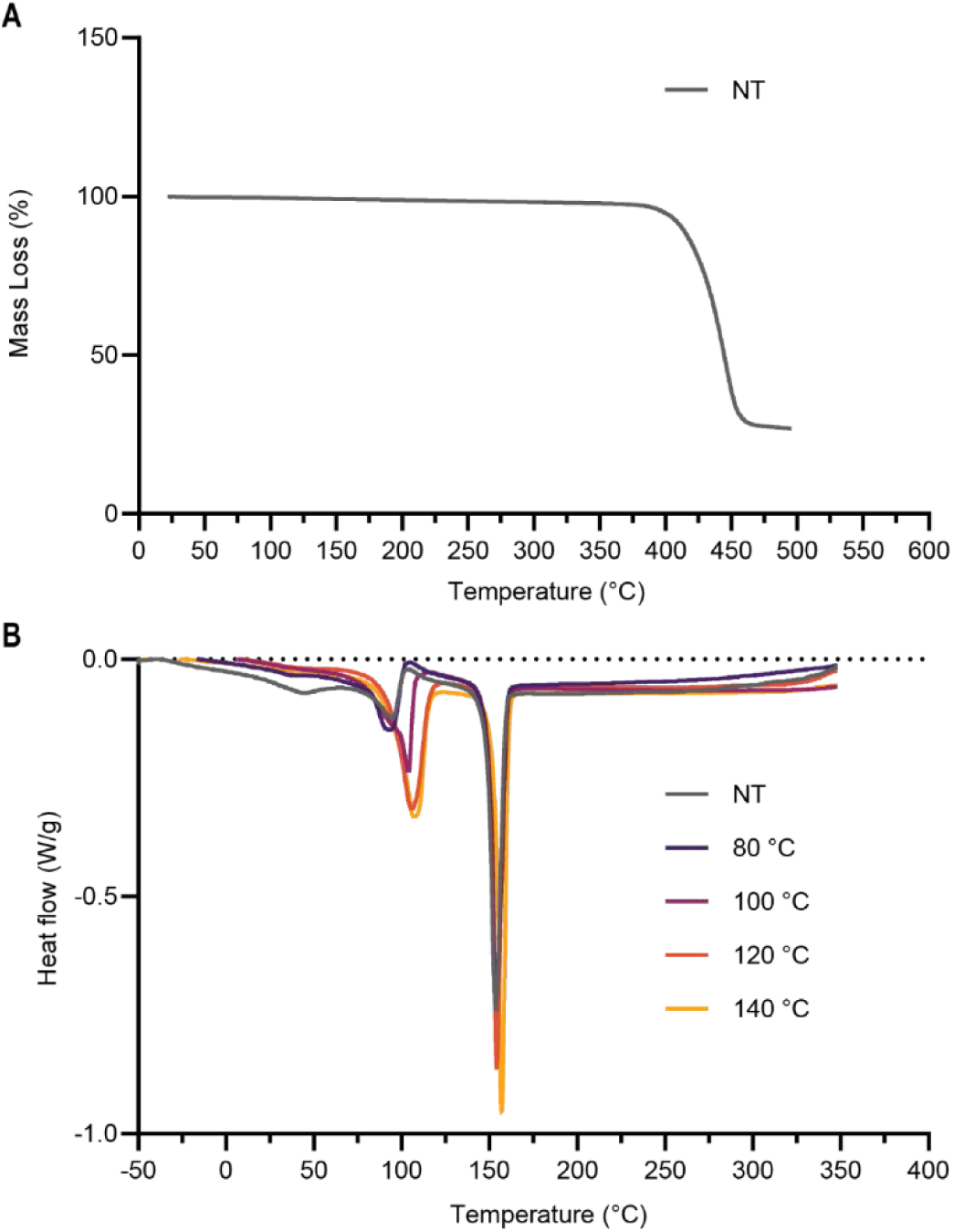
Thermal analysis of PVDF-TrFE. A) Thermogravimetric analysis (TGA) graph of NT PVDF-TrFE showing a percentage of initial mass lost as sample is heated. B) Differential scanning calorimetry (DSC) graph of NT PVDF-TrFE nanofibers and annealed PVDF-TrFE at 80°C, 100°C, 120°C and 140°C to examine thermal transitions and overall crystallinity of the fibers.

Total crystallinity fraction (%) is shown to steadily increase with rising temperature. Specifically, crystallinity fraction (%) increased from 15.43 to 29.39 % in the NT to the 140°C groups respectively, indicating a 90.4 % increase. The total crystallinity fraction of PVDF-TrFE annealed scaffolds is highlighted in Table S1.

### 2.2 Morphological properties

We conducted scanning electron microscopy (SEM) to observe changes in the morphology of annealed PVDF-TrFE nanofibers across a range of annealing temperatures (80°C, 100°C, 120°C, and 140°C). SEM images obtained at different temperatures facilitated the assessment of alterations in fiber morphology (Figure 2). As annealing temperature increases, the presence of lamellar-like structures becomes increasingly evident. 2D Fourier image processing technique was used to validate these morphological changes. This method produced polar plots displaying needle-like structures with singular alignment, while branched polar plots showed strong alignment across a wider range of directions, indicating the presence of lamellar-like structures [53].

**Figure 2:**
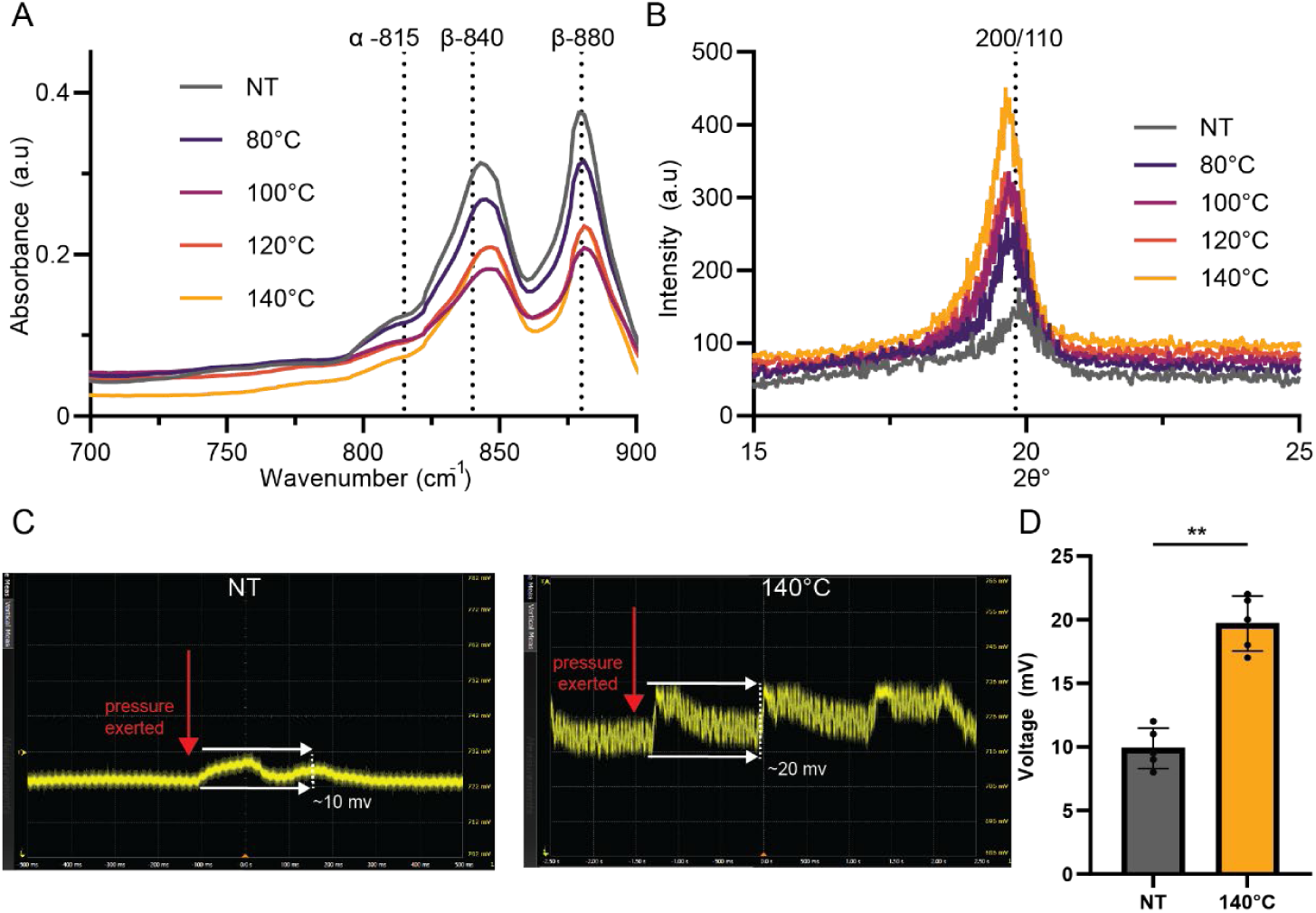
Morphological Assessment of annealed PVDF-TrFE nanofibers. A-E left) SEM images of annealed nanofibers NT, 80°C, 100°C, 120°C and 140°C respectively (scale bar = 500 microns). A-E right) shows respective polar plots indicating fiber orientation.

SEM at a high magnification of 8000x was used in order to quantify and analyze fiber diameter, alignment, and porosity (Figure 3). The investigation revealed a relatively uniform fiber diameter across the annealed groups. The scaffold group annealed at 140°C exhibited the smallest average fiber diameter. However, variations in fiber diameter were likely due to batch-to-batch differences rather than the annealing process itself, as the melting temperature was not exceeded. A detailed summary of these findings is provided in Table S2. An interesting trend emerged regarding porosity calculated (Eq 2), where *Image Area_Total_* is the total area of the SEM image and *Image Area_Fibers_* is the total area of the SEM image covered by PVDF-TrFE nanofibers.

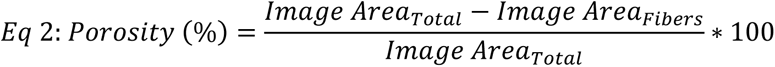

**Figure 3:**
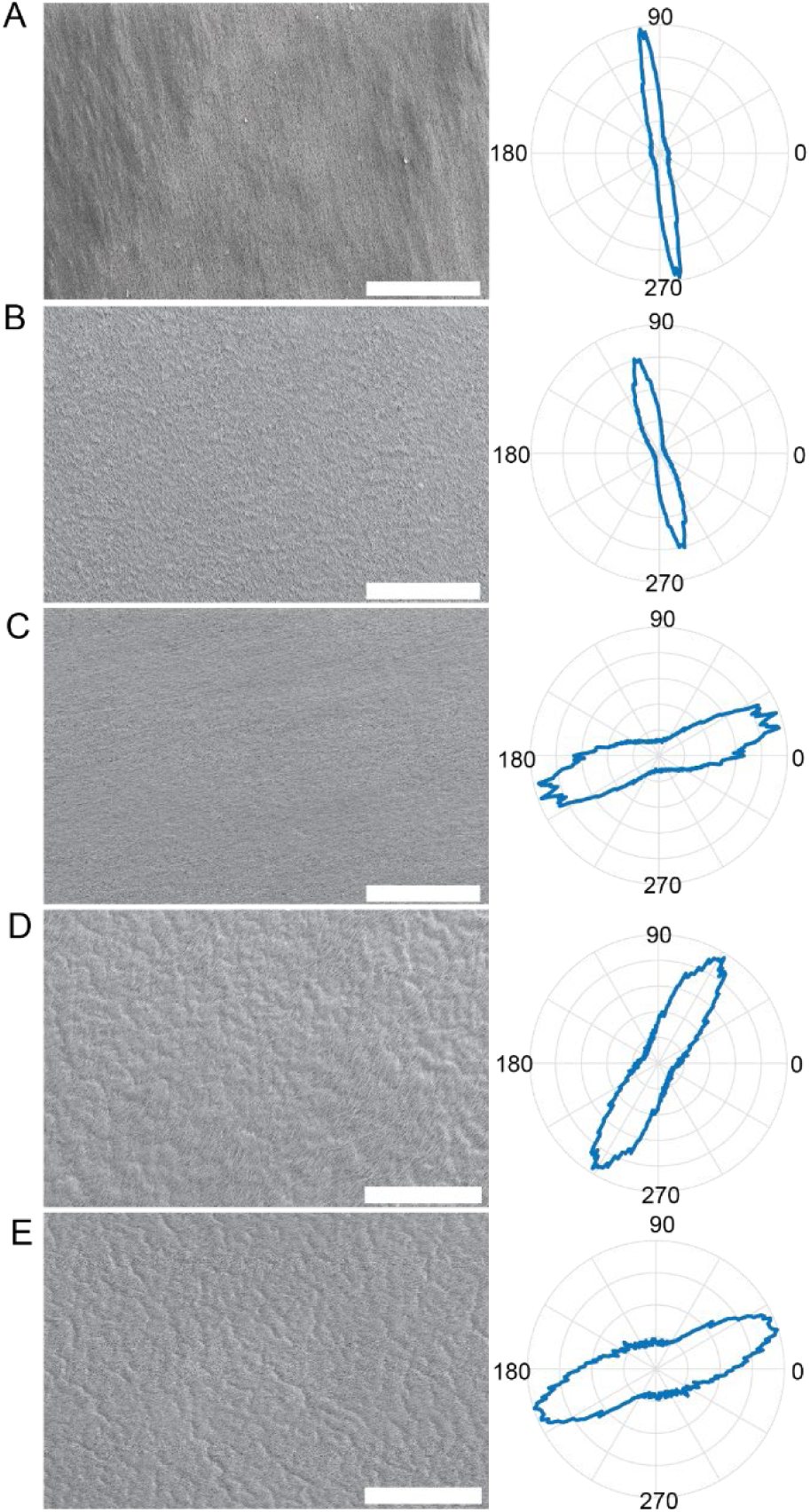
Scanning electron micrographs and properties. A-E left) SEM images of nanofiber groups NT and scaffolds annealed at 80°C, 100°C, 120°C, and 140°C. (scale bar = 5 micrometer). A-E) right shows respective fiber diameter histograms. F) Alignment Index and G) porosity of annealed fibers with results reported as the mean ± SD of five independent experiments (***p<.001).

As the annealing temperature increased, so did the porosity of the scaffolds in (Figure 3G). However, significant variations in nanofiber alignment were predominantly observed between the groups annealed at 80°C and 140°C (Figure 3F). Despite these differences, a high level of fiber alignment was consistently maintained across all experimental groups, underscoring the effectiveness of the annealing process in preserving morphological characteristics while modifying microstructural properties.

### 2.3 Mechanical Properties

Tensile testing to measure Young’s modulus showed distinct stiffness variations across treatment groups. The NT group had a Young’s modulus of 11.2 MPa. The 80°C annealed group had the lowest stiffness at 1.14 MPa, while the 140 °C annealed group showed the highest at 71.8 MPa (Figure 4A). Ultimate tensile strength assessments revealed that the NT group had a strength of 11.23 MPa, with the 140°C group achieving the highest strength of 17.8 MPa (Figure 4B). Additional mechanical properties such as substrate stiffness, were assessed using atomic force microscopy (AFM), where the NT group’s modulus was 56.1 kPa. The 120°C group exhibited the highest modulus at 163 kPa, closely followed by the 140°C group at 160 kPa, showing negligible differences (Figure 4C). These results indicate a clear trend of increased stiffness with higher annealing temperature, underscoring the impact of annealing on the mechanical properties of the scaffolds, with values reported in Table S3.

**Figure 4:**
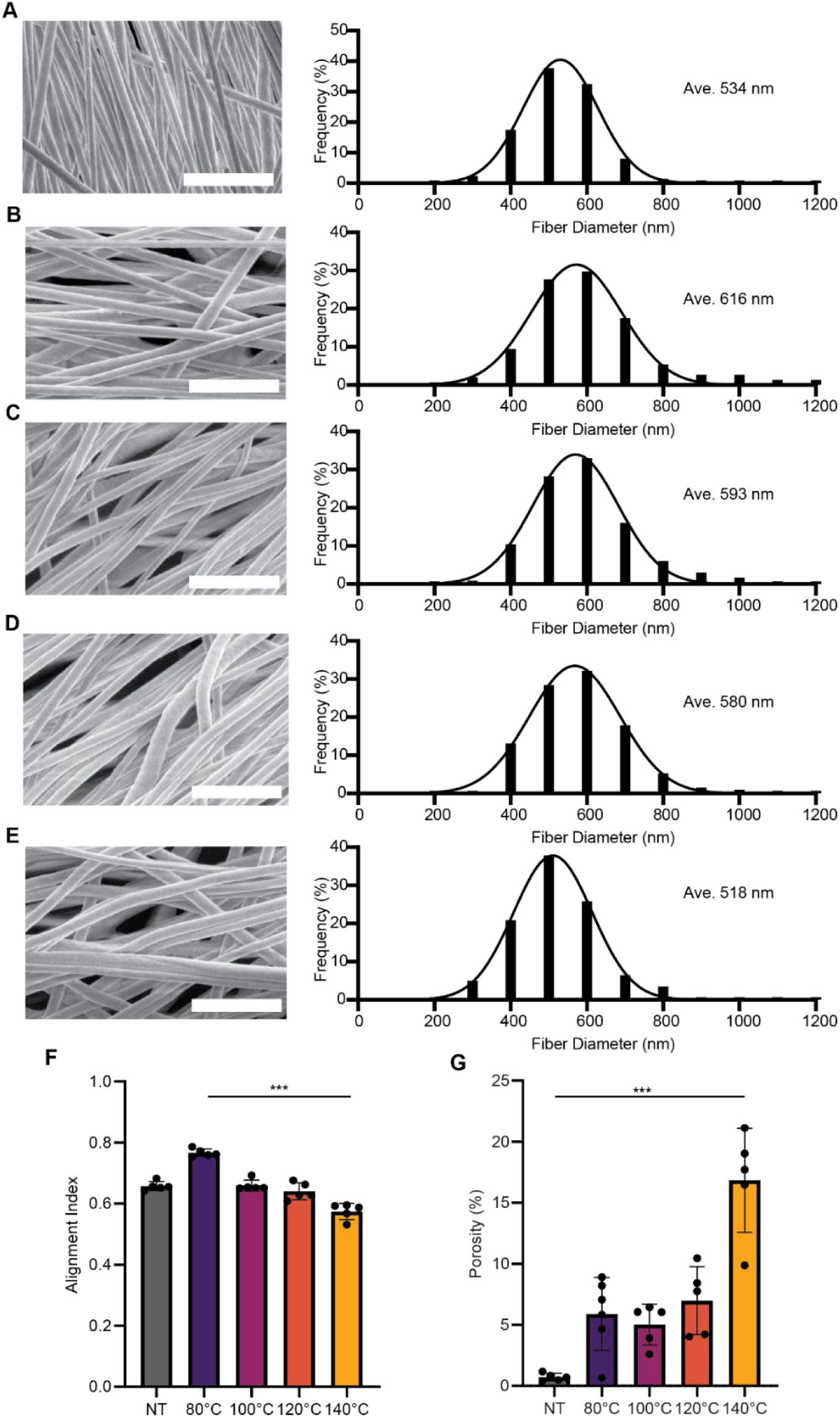
Mechanical properties of annealed PVDF-TrFE. A) Bulk Young’s Modulus of annealed PVDF-TrFE nanofibers undergoing longitudinal tensile testing in direction of alignment with results presented as the mean ± SD of four independent experiments (∗p<0.05, **p<.01). B) Ultimate Tensile Strength of annealed PVDF-TrFE nanofibers with results presented as the mean ± SD of four independent experiments (∗p<0.05, **p<.01). C) Atomic Force Microscopy of annealed on PVDF-TrFE to indicate stiffness on surface of nanofibers results presented as the mean ± SD of a hundred independent indentation values (****p<.0001). Not all significances shown in figure, reported in Table S4.

### 2.4 Piezoelectric Properties

To investigate the piezoelectric properties of NT and annealed scaffolds, Fourier transform infrared spectroscopy (FTIR) and X-ray diffraction (XRD) were utilized. Due to the semi crystalline nature of PVDF-TrFE, it is necessary to quantify to which extent the amorphous α-phase of the scaffold transitions to the β-phase crystallinity annealing using FTIR [54]. FTIR reveals characteristic β-phase bands of PVDF-TrFE at 840 cm⁻¹ and 880 cm⁻¹, as well as an α-peak band at 766 cm⁻¹ (Figure 5A). The ratio of absorbance values is used to calculate the corresponding to the F(β) crystallinity fraction through the Lambert Beer Law (Eq 3), where Aβ and Aα are the absorbances at 840 cm^−1^ and 766 cm^−1^, respectively. Kα and Kβ represent the absorption coefficients at the respective wavenumbers, which are 6.1 × 10^4^ cm^2^/mol and 7.7 × 10^4^ cm^2^/mol [55, 56].

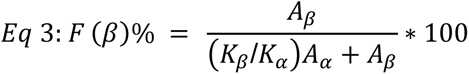

**Figure 5:**
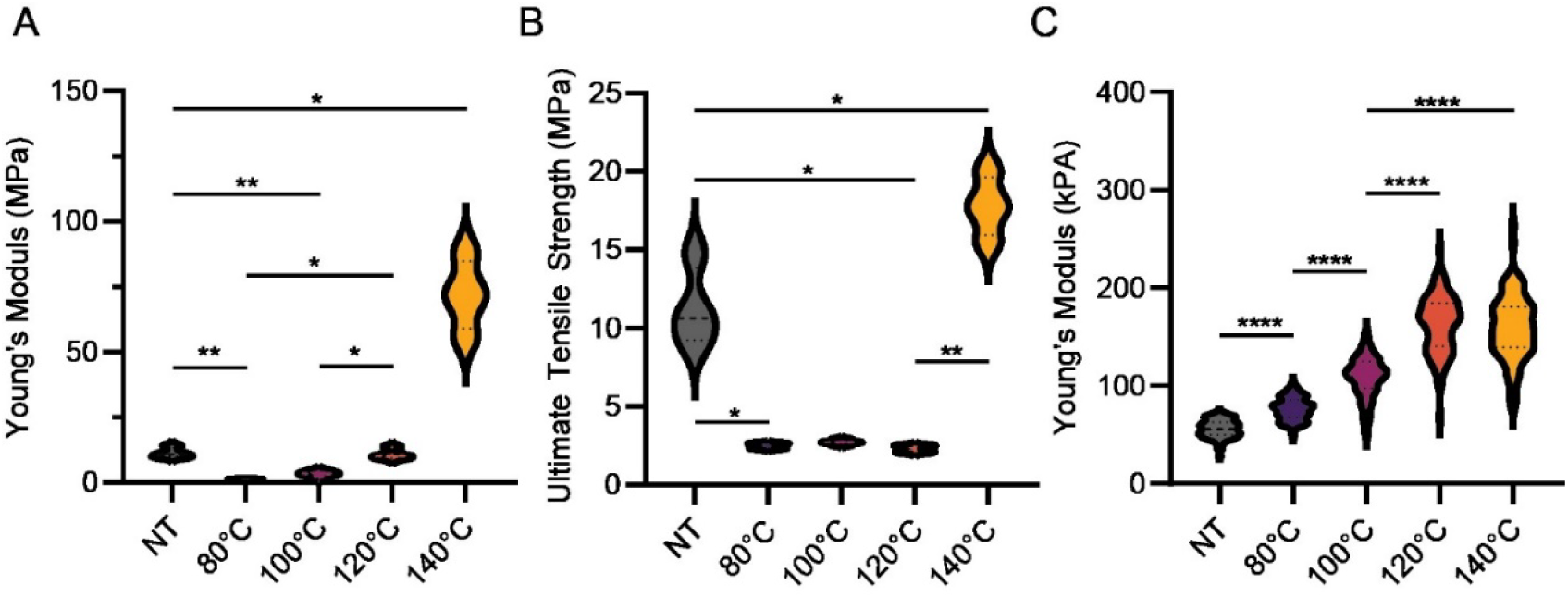
Piezoelectric responses of annealed PVDF-TrFE. A) Fourier transform infrared spectroscopy of respective annealed and unannealed PVDF-TrFE groups. B) X-ray diffraction of groups C) Oscilloscope unfiltered electrical output from piezoelectric response in NT (left) and 140°C (right) when stimulated by tension corresponding to consistent voltage readouts. D) Bar plot of filtered maximum peak-to-peak voltage outputs of NT and 140°C annealed PVDF-TrFE scaffolds (n=5) when subject to deformations (**p<.01).

The F(β) percentages decreased from the NT group to the annealed groups at 80°C and 100°C. However, at higher annealing temperatures of 120°C and 140°C F(β) percentages increased with the annealed 140°C group having higher than NT the respective values shown in Table S5. The XRD patterns for the annealed PVDF-TrFE scaffolds displayed a pronounced diffraction peak at 2θ = 19.8°, indicative of the 110/200 reflection of the orthorhombic β-phase crystal across all groups [57]. Notably, the 140°C annealed group manifested the most pronounced β-phase peak relative to the other groups, signifying highest β phase percentage of crystallinity (Figure 5B). To measure the maximum electrical output of the piezoelectric scaffolds, NT and 140°C annealed scaffolds were fabricated into chips using a cantilever setup as depicted in our previous publications [30, 31]. These scaffolds were then deformed to generate consistent electrical signals, and the results were recorded using an oscilloscope indicating an approximate two-fold increase in the 140°C group compared to NT (Figure 5C-D).

### 2.5 In Vitro Biocompatibility and Cellular Responses

To investigate the responses of cells cultured on annealed scaffolds, Schwann cells (SCs) and fibroblasts at a density of 100 cells per mm² were cultured on both NT and 140°C annealed scaffolds. Initial observations revealed comparable levels of cell adherence on both substrates within the 72-hours (Figure 6A). Analysis was employed to evaluate cellular viability and proliferation, using immunofluorescence of SCs on both scaffold groups with MKI67 marker, a well-established indicator of cellular proliferation (Figure 6A). Metabolic activities of SCs on both substrates were assessed by 3-(4,5- dimethylthiazol-2-yl)-2,5-diphenyl-2H-tetrazolium bromide (MTT) assays over the first 48 hours of incubation. Both the NT and 140°C groups showed no significant difference in supporting SCs (Figure 6B) and fibroblast viability (Figure S1). Using DAPI staining for total cell count, facilitated the quantification of proliferating SCs. The proliferation rates of SCs were determined to be 5.9±1.2% for the NT group and 8.3±2.4% for the 140°C annealed group, indicating a notable increase in the proportion of actively proliferating cells in the 140°C group by approximately 40%, relative to the NT group (Figure 6C). Western blot analysis of SC lysates 48 hours post-seeding on scaffolds revealed an increase in RhoA and FAK cytoskeletal protein and cell to matrix adhesion protein respectively, along with a decrease in the cell-to-cell adhesion protein N-cadherin in the 140°C annealed scaffold group compared to the NT group (Figure 6D) with raw Western blots depicted (Figure S2).

**Figure 6:**
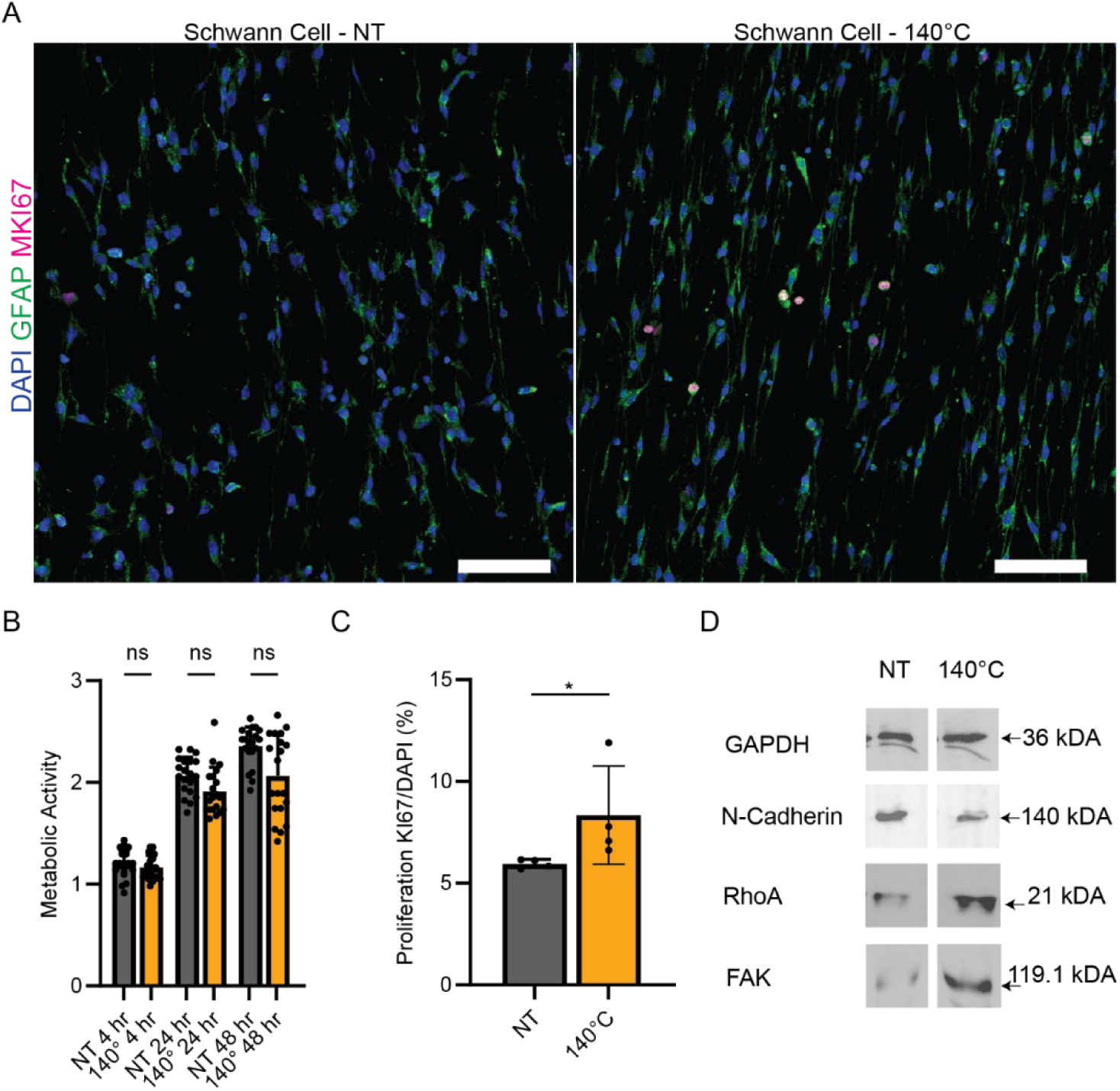
Cellular responses to the annealed PVDF-TrFE. A) Representative images showing SCs after 48 hours with DAPI (blue), GFAP (green), and MKI67+ (red) for NT (left) and 140°C (right) (scale bar 100 microns). B) Bar graph of MTT cell metabolism assay comparing SC viability on NT and 140°C groups over 48 hours (n > 12 readings). C) Bar graph of Schwann cell proliferation using MKI67+/DAPI (%) between NT and 140°C groups (n = 4) (∗p < 0.05). D) Western Blots of GAPDH (control), N-Cadherin (cell-cell attachment protein), RhoA (cytoskeletal protein), and FAK (focal adhesion kinase) using lysates from SCs seeded on NT (left) and 140°C (right) after 48 hours.

### 2.6 RNA-Sequencing

SCs were cultured on NT and 140°C PVDF-TrFE scaffolds for 72 hours, after which RNA quality was assessed (Figure S3). Resultant RNA was sequenced utilizing the Rattus_norvegicus (UCSC rn5) reference genome with results visualized (Figure 7). PCA analysis revealed that SC RNA from the NT and 140°C groups formed two distinct clusters (Figure 7A). Differential gene expression analysis identified a total of 300 differentially expressed genes, with 187 genes upregulated and 113 genes downregulated when comparing RNA from the baseline NT group to 140 °C group (Figure 7B). Notably, genes associated with extracellular matrix (ECM) production, such as Adamts1, Agrn, Loxl2, P4ha2, P4ha3, Serpinb6a, and Tgfb1i1, were significantly upregulated [58, 59]. Additionally, there was upregulation of collagen genes (Col18a1, Col4a1, Col5a2) [60, 61]. Collagens are the predominant structural proteins in the ECM, providing tensile strength and playing crucial roles in cell adhesion, migration, and proliferation [62]. Glycoprotein genes (Igfbp5, Igfbp7) were also upregulated [61]. Glycoproteins in the ECM and basement membrane are essential for cell recognition, adhesion, migration, and proliferation. These ECM remodeling genes are important for maintaining and restructuring the ECM, thereby promoting a regenerative microenvironment [63]. Genes coding for growth factor related proteins (CTGF, CYR61, ESM1, GDF15, PDGFB, PGF) were significantly upregulated, highlighting their roles in promoting cell adhesion, migration, proliferation, and angiogenesis [64–67]. Upregulation of these genes enhances vascularization and blood supply, thereby facilitating tissue repair and regeneration. This highlights the scaffold’s efficacy in creating a regenerative microenvironment that promotes engraftment [68–70]. Additionally, genes involved in cytoskeletal dynamics (Actn1, Actr1b, Csrnp1, Csrp2, Dact3, Epha2, Fkbp11, L1cam, Pdlim1, Pdlim7, Rhob) were upregulated, highlighting their roles in cell adhesion, migration, and signaling processes crucial for tissue repair [58, 59, 61, 71, 72]. The cytoskeleton orchestrates dynamic processes like wound repair by forming contractile arrays and cellular protrusions. These processes are driven by actin, myosin, and microtubule networks, and regulated by Rho family GTPases [64].The upregulation of ion channel genes (Atp2b4, Aqp1, Cyba, Gpx8, Slc2a3, Slc39a7, Slc6a6, Tspo) in SCs indicates enhanced intracellular signaling and ion transport, which are essential for maintaining cellular homeostasis [73–75]. Although not fully explored in the context of tissue repair, this upregulation suggests increased metabolic activity and energy levels in SCs, promoting their proliferation and function in nerve regeneration [76]. The upregulation of calcium signaling genes (Cald1, S100a10, S100a4, S100a6, Anxa5, Tpm1) is related to jumpstarting the regenerative cascade and promoting cellular migration as well as proliferation. Interestingly genes related to proteins involved in peripheral nerve repair cascade, regulating fibrosis, and relating to angiogenesis were upregulated (Creb3l1, Ecm1, Ier3, mdm2, Mvp, myo1c, Ngfr, Nrp2, and Prdx5) with several downregulated genes involved in myelination and cell-cell adhesions (Chd6, Mest, and Mpz) [77–82]. DEGs of interest are depicted in heat map (Figure 7C).

**Figure 7:**
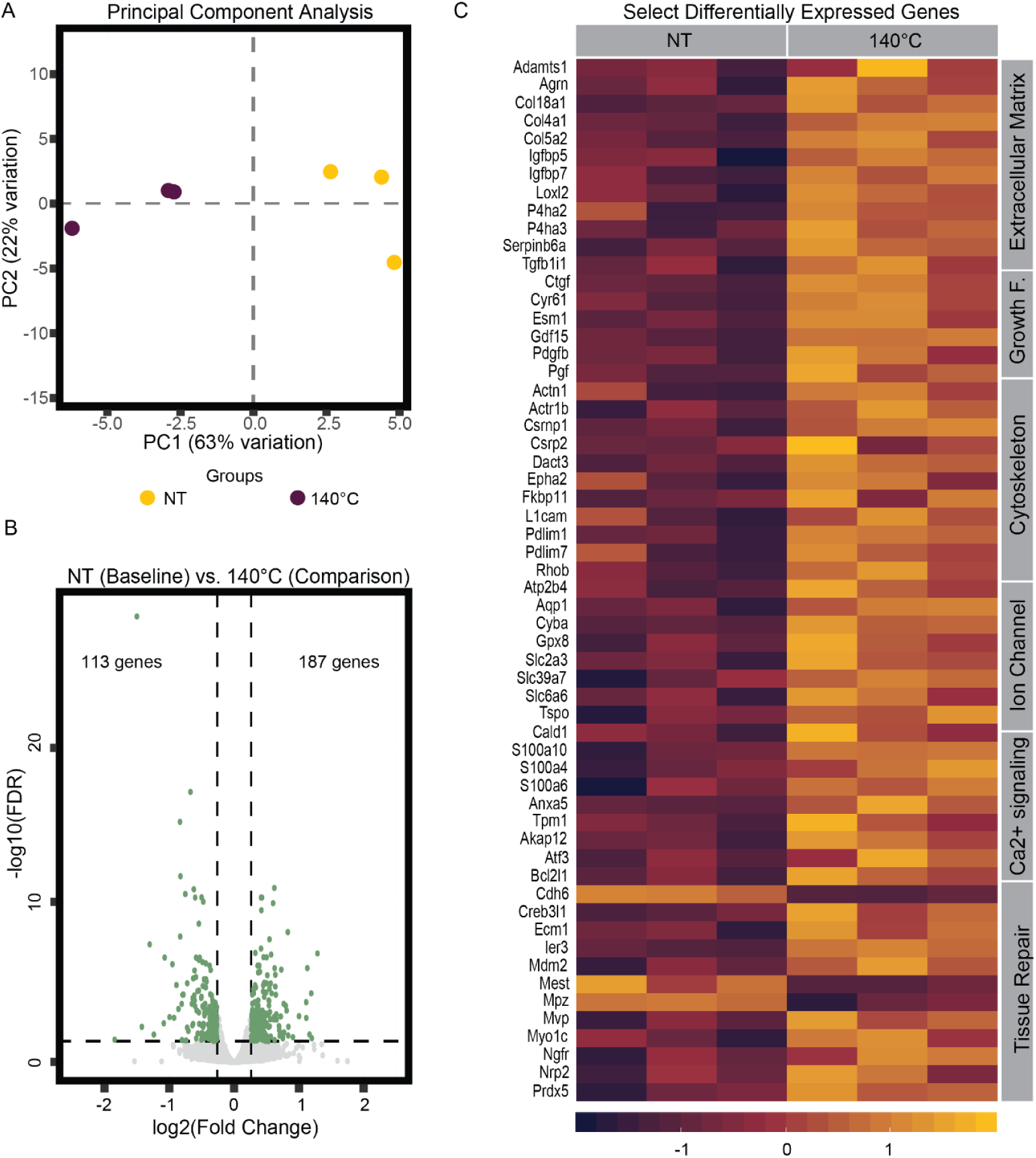
Differential Gene Expression of SCs on NT PVDF-TrFE scaffolds and annealed PVDF-TrFE scaffolds at 140°C. A) Principal component analysis (PCA) illustrates the clustering of sample groups, showing that both NT and annealed groups form distinct clusters. PC1 accounts for 63% of the variation, while PC2 explains 22% of the variation. B) Volcano plot displays the differentially expressed genes (DEGs) with a total of 300 genes highlighted in green out of 9580 assessed. Among these, 187 genes are upregulated, and 113 genes are downregulated in the annealed group compared to the NT baseline. DEGs were identified using an absolute log2 fold-change threshold of 0.26303 (equivalent to a fold-change of 1.2). C) Heatmap of selected upregulated DEGs in Schwann cells cultured on PVDF-TrFE scaffolds annealed at 140°C. The heatmap shows gene expression levels ranging from highly upregulated (yellow) to highly downregulated (dark purple). Genes are listed row-wise, and sample groups are displayed column-wise (n=3 per group). A grey column on the right categorizes the genes into functional groups: Extracellular Matrix, Growth Factors, Cytoskeleton, Ion Channels, Calcium (Ca²⁺) Signaling, and Tissue Repair.

### 2.7 In Vivo Foreign Body Response Assessment

To assess foreign body response, NT and 140 °C annealed PVDF-TrFE scaffolds were subcutaneously implanted ventrally into C57BL/6 mice, interacting with stomach muscle and skin layers, and stitched up (Figure 8A). We examined tissue responses to these nanofiber materials four weeks after implantation by analyzing 24 slides stained with hematoxylin and eosin (H&E) and Sirius Red (SR). The slides were evaluated in a blinded analysis by a pathologist (Figure 8B). Results consistently revealed linear, polarizable nanofibers beneath the skeletal muscle without epithelial encapsulation. Both nanofiber groups triggered a response featuring multinucleated giant cells and connective tissue fibrosis but did not form a distinct capsule four weeks post implantation. This inflammatory response, marked by multinucleated giant cells, neutrophils, and fibrosis, was consistent, demonstrating that tissue reactions to NT and 140°C annealed scaffolds were undifferentiable.

**Figure 8:**
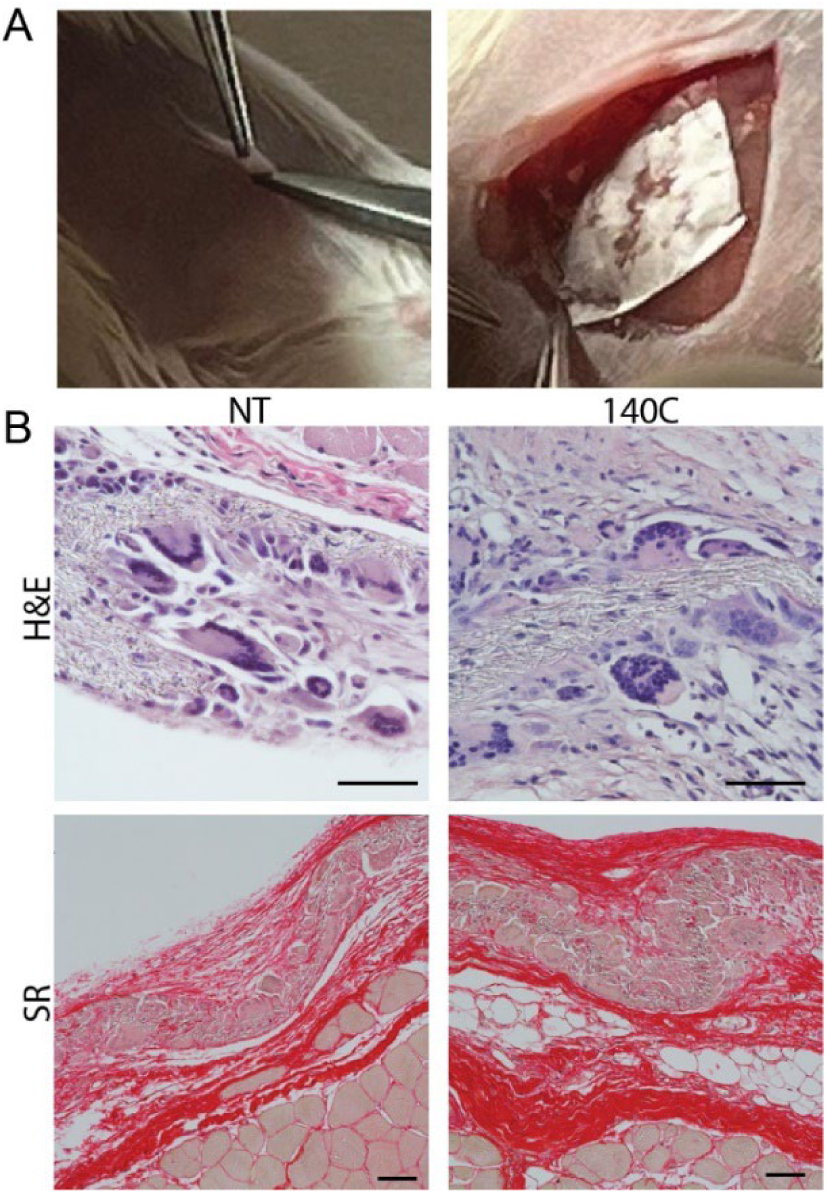
In vivo foreign body responses to annealed scaffolds. A) Surgical implantation of annealed PVDF-TrFE scaffolds into mice. B) Representative images showing Hematoxylin and eosin (H&E) staining with picrosirius red staining post 28 days after implantation of NT & 140 PVDF-TrFE scaffold groups (n=4 per group) in C57BL/6 mice with scale bar = 50 microns.

## 3. Discussion

This study addresses the challenge of low electrical responses in soft piezopolymers, compared to composite piezoelectric counterparts that incorporate metallic ions or conductive elements. We systemically investigate the effects of annealing temperatures on PVDF-TrFE to enhance piezoelectric properties. Thermal analysis using DSC revealed that annealing PVDF-TrFE scaffolds at 140°C optimizes their ferroelectric and piezoelectric properties. DSC analysis demonstrated a continual increase in total crystalline content and β-phase crystallinity, reaching a 90% improvement at 140 °C. Higher crystallinity in the β-phase, confirmed through FTIR using Lambert-Beer’s Law, is directly linked to improved piezoelectric sensitivity. XRD analysis further supported this, showing the highest relative intensity of the 200/110 peaks in the 140°C group [83, 84]. These findings are consistent with other studies showing that annealing PVDF-TrFE sputter-coated films at similar temperatures enhances beta-phase crystallinity, thereby improving energy harvesting and related applications [85, 86]. PVDF-TrFE electrospun nanofibers differ from films in their superior high spatial resolution and ECM like morphology making them more translatable as tissue engineered grafts [84]. Annealing PVDF-TrFE resulted in enhanced porosity and lamellar like structures, indicating enhanced biomaterial suitability by providing space for cell penetration and nutrient diffusion in the 140°C group [87].

Annealing at 140°C significantly improved the mechanical properties of PVDF-TrFE scaffolds, resulting in approximately six-fold increase in tensile Young’s modulus and a 60% increase in ultimate strength. The substrate stiffness, measured by AFM, increased three-fold in the 140°C group annealed group compared to NT, provides insights into the mechanical environment experienced by cells and tissues at the surface level [88, 89]. This enhancement is expected, as annealing increases stiffness and ultimate strength in certain polymers by enhancing crystallinity, reducing defects, and improving molecular orientation [44, 86, 90, 91]. Consequently, as the load-bearing capacity increases, the scaffolds can handle higher mechanical forces and are able to generate a larger piezoelectric response. Favorably, the enhanced substrate stiffness (50-160 kPa) falls within the range of natural soft tissues, making these scaffolds suitable for mimicking the mechanical environment of tissues like nerves, skin, and arteries [92]. Interestingly, while AFM showed increased substrate stiffness with higher annealing temperatures, Young’s modulus and load-bearing strength drastically plummeted when scaffolds were annealed at temperatures of 80°C, 100°C, and 120°C. This decline in mechanical strength is likely due to inconsistent crystalline regions throughout the scaffolds causing higher frequency of tears under tension, a phenomenon not previously reported for annealed PVDF-TrFE [41, 93–95].

The piezoelectric peak to peak electrical signals of annealed scaffolds were significantly improved, with the 140°C annealed scaffolds generating an electric signal twice of NT scaffolds. Although, piezoelectric coefficient (d33) was not directly measured in this study, we expect there to be an increase of approximately 30-40% in d33 coefficient as seen when PVDF electrospun fibers were annealed with similar treatment and duration at 140°C [96]. In this case controlled mechanical stimuli, such as shockwaves or ultrasound, enable the annealed scaffold to produce a 30-40% improved electrical response, making it more effective in soft tissue repair where external stimulation is beneficial [97, 98]. Murillo et al. identified that local electric fields generated by cell traction forces during cell-material interactions can modulate cell behavior on the piezoelectric substrate [99]. The piezoelectric response is influenced by the electromechanical properties of the material and the cell’s adhesion state, which determines the location and magnitude of the applied cellular traction forces [99, 100]. Xie et al. further modeled this phenomenon in COMSOL, showing that aligned electrospun PVDF nanofibers produced a voltage of -21 mV under an approximate force of 10 nN force, compared to 18.9 mV for randomly oriented fibers verifying with measuring intracellular calcium levels [101, 102]. Annealing PVDF-TrFE scaffolds at 140°C maximizes their energy harvesting potential, allowing them to generate electrical stimulation signals in the range of endogenous-like electrical fields (20 – 300 mV/m) similar to those observed in human skin wounds under similar force and deformation estimates of nanofibers [103–106]. Thus, we expect PVDF-TrFE scaffolds annealed at 140°C to produce an even higher level of electrical response compared to PVDF and NT groups under similar activation forces.

Given that substrate stiffness directly correlates with increased cellular traction forces and focal adhesions, we investigated whether the 140°C annealed scaffold could promote phenotypical changes in cells [107, 108]. We hypothesized that the increased cellular traction forces and additional points of contact on the stiffer, more piezoelectric polymer would amplify the effect of electrical stimulation and promote advantageous cellular and tissue repair responses. This amplification would vary depending on cellular movement and focal adhesions, potentially enhancing the piezoelectric forces that stimulate the cells. To verify this, we first confirmed that SCs seeded on 140°C annealed PVDF-TrFE scaffolds exhibited comparable metabolic activity compared to NT scaffolds and showed increased proliferation, as indicated by the KI67 marker in the 140°C group. Western blot analysis revealed that protein levels of focal adhesion kinases (FAK) and RhoA, which are directly linked to cellular traction forces and adhesion to the matrix, were elevated in the 140°C group, while N-cadherin, which mediates cell-cell adhesions, was decreased. RNA-sequencing of SCs on NT and 140°C annealed PVDF-TrFE scaffolds further demonstrated that genes coding for ECM proteins and cytoskeletal dynamics creating a favorable microenvironment tissue engraftment and repair [87, 109–115]. Additionally, genes that code for protein growth factors such as connective tissue growth factor (CTGF) and placental growth factor (PGF) genes were upregulated thus promoting angiogenic factors that aid in vascularization and incorporation of the graft into the tissue [69, 116, 117]. Genes associated with ion, transmembrane channels and calcium signaling, including those from the S100 protein family and other calcium dependent genes like Cald1, Anxa5, and Tpm1, were upregulated. These findings align with a study that showed further electrical stimulation could lead to a substantial increase in the expression levels of S100 protein related genes and the nerve growth factor receptor (Ngfr) gene, which are associated with myelination in SCs and an increase in nerve growth factor production, both critical for peripheral nerve repair [118–120]. This suggests that the annealed group had higher electrical stimulatory effect of the scaffold on SC phenotype due to increased net electrical charges from cellular traction forces, resulting in elevated ion flux and intracellular calcium changes, providing evidence of enhanced scaffold-induced cellular responses.

One of the biggest limitations of allografts and various implantable biomaterials is foreign body response that can illicit high levels of inflammatory immune activity to hinder tissue regeneration in all tissue types [121–123]. We found that the consistent tissue response in C57BL/6 mice models, characterized by the presence of multinucleated giant cells and connective tissue fibrosis without distinct capsule formation, suggests that the nanofiber scaffolds, regardless of annealing treatment, elicit a manageable inflammatory response. This uniform response across both PVDF-TrFE scaffold groups underscores the potential of these materials for acute tissue repair in clinical settings. Similarly, a recent study has shown that piezoelectric polyurethane (PU)/PVDF electrospun scaffolds implanted subcutaneously had a foreign-body reaction with minimal chronic inflammation after fourteen days for wound healing applications in Sprague Dawley rats [124, 125]. Another study evaluated the in vivo biocompatibility of a PTFE-PVDF-PP terpolymer membrane for glaucoma treatment in rabbits using the non-perforating deep sclerectomy procedure. It found a similar slight fibrotic response in both the control and no-implant groups after one month, indicating that fibrosis levels remain low within this period for implants intended for longer use [126]. In essence, we demonstrate that PVDF-TrFE scaffolds annealed at 140°C can be implemented subcutaneously. However, further research is needed to address functional recovery in targeted physiological tissue systems, such as nerve and skin. Future studies should evaluate the long-term effects of PVDF-TrFE as a foreign body, both subcutaneously and within specific regenerative contexts, with and without external activation through mechanotherapy. This will help to better understand the tissue-material interactions and their impact on inflammation when activated as well as specific tissue repair outcomes.

## 4. Conclusion

Our investigation into the effects of annealing on electrospun PVDF-TrFE scaffolds has identified significant enhancements in biomaterial properties, aimed at improving bioelectrical stimulation capacity for tissue repair. By annealing scaffolds at 80°C, 100°C, 120°C, and 140°C, we determined that 140°C optimizes mechanical properties, porosity, and beta phase crystallinity, as confirmed by XRD, FTIR, and DSC analyses. PVDF-TrFE scaffolds annealed at 140°C resulted in a two-fold increase in electrical output under mechanical stimulation compared to NT scaffolds. Both NT and 140°C annealed scaffolds exhibited comparable biocompatibility with fibroblasts and Schwann cells. Notably, Schwann cells showed higher proliferation on the 140°C annealed scaffolds using a KI67 marker. Western blot analysis revealed elevated levels of FAK and RhoA proteins, which are linked to cellular traction forces and cell-matrix adhesion, in the 140°C annealed group. RNA-sequencing of Schwann cells on the 140°C annealed scaffolds demonstrated upregulation of genes involved in extracellular matrix components, cytoskeletal dynamics, growth factors, ion channels, and calcium signaling, indicating an improved microenvironment and enhanced electrical stimulation conducive to a regenerative phenotype. In vivo studies indicated that both NT and 140 °C annealed scaffolds exhibited similar polymeric-tissue responses, characterized by the presence of giant cells but no distinct rejection after 4 weeks of implantation. These findings support the potential of 140°C annealed PVDF-TrFE scaffolds for advanced regenerative therapies, providing enhanced electrical stimulation and improved functional recovery in tissue regenerative graft applications.

## 5. Materials & Methods

### 5.1 Materials

N,N-Dimethylformamide (Sigma-Aldrich, 68-12-2) and acetone (Sigma-Aldrich, 67-64-1) were used as solvents used in electrospinning. Piezotech P(VDF-TrFE) copolymer resin 70/30 (PolyK-Technologies) was used for electrospinning, and conductive silver paint (Ted Pella Inc) was utilized for electrical connections. Phosphate Buffered Saline (PBS) (Corning, 21-040-CM) was used for cell culture procedures, along with bovine serum (Gibco, 12105) and fetal bovine serum (FBS) (Gibco). The MTT reagent (3-(4,5- dimethylthiazol-2-yl)-2,5-diphenyltetrazolium bromide) (MilliporeSigma, 298-93-1) was employed for cell viability assays. For Western blotting, N-Cadherin Antibody (13A9), RhoA Antibody (1A11-4G10), and FAK Antibody (OTI4D11) were used (NovusBio). Mouse anti-GFAP (Thermo-Fisher, MA5-12023) and rabbit anti-MKI67 (Thermo-Fisher, RM- 9106) antibodies were utilized for immunohistochemical staining. Isoflurane (Butler Schein, 66794-017-25) and Buprenex (Midwest Veterinary Supply) were used for anesthesia and analgesia, respectively. Pen/Strep and Amphotericin B (Midwest Veterinary Supply) were used to prevent contamination in cell cultures. Paraformaldehyde (Sigma-Aldrich) was used for tissue fixation, and histological staining was performed using a Picrosirius red kit (Polysciences Inc, 24901-500) and Hematoxylin and Eosin (H&E) (Sigma-Aldrich, 1.15973). For RNA sequencing, total RNA was isolated using the mirVana kit (Thermo Fisher, Waltham, MA). The quality of the total RNA was analyzed using a Bioanalyzer (Agilent, Santa Clara, CA).

### 5.2 Preparation of Nanofiber scaffolds

PVDF-TrFE nanofibers were prepared as described previously by our group [30]. Briefly, a polymer solution consisting of 20 (w/v) % PVDF-TrFE (70/30) and solvent of DMF and acetone (6/4 v/v) was added to a 5 mL syringe fitted with a 20-gauge needle. The syringe pump was set to supply a flow rate of 1 mL h^−1^. The needle tip was positioned 10 cm away from the collector, which rotated at 2000 RPM to produce aligned fibers wrapped on a conductive aluminum foil sheet. An 18 kV voltage was applied between the needle tip and the collector. The electrospinning process is run for three hours.

### 5.3 Annealing of PVDF-TrFE scaffolds

Scaffolds were apportioned and subjected to heat in a thermally controlled convection oven (Lindberg Box Furnace, Fisher Scientific) for 90 minutes at temperatures of 80°C, 100°C, 120°C, and 140°C, respectively. Following this heat treatment, the scaffolds were promptly removed and rapidly cooled using 100% isopropyl alcohol at -20°C. The scaffolds were then left to air dry for 24 hours before being utilized.

### 5.4 Thermal Characterization of PVDF-TrFE and total Crystallinity

Thermal properties of PVDF-TrFE scaffolds were assessed using Thermogravimetric Analysis (TGA) and Differential Scanning Calorimetry (DSC). TGA, performed with TGA 550 (TA Instruments), evaluated weight changes in a 3.683 mg scaffold sample, heated at 5.00°C/min to 500.00°C/min, to record the mass loss. DSC analysis, conducted with DSC D2500 (TA Instruments), measured energy exchange during heating and cooling.

5.50 mg samples of PVDF-TrFE scaffolds were annealed at systemic annealing temperatures and were analyzed over -50°C to 350°C at a rate of 10°C/min, assessing melting, and overall crystallinity behaviors to calculate overall crystallinity for scaffold groups.

### 5.5 Scanning Electron Microscopy and Image Analysis

Scanning electron microscopy (SEM) (ApreoC SEM, ThermoFisher) was used to analyze the morphology of electrospun PVDF-TrFE nanofibers annealed at different temperatures. Three samples per experimental condition were prepared from electrospun scaffolds, and were sputter coated with a layer of gold for approximately 10 seconds. Five images per sample were taken with a 5 mm working distance and an acceleration voltage of 2 kV. SEM images were captured with an FEI XL-30 microscope (Low-Vac) using an EDAX elemental analysis detector. Resultant scaffold images (n=5) were analyzed using ImageJ software (version 1.53k) for porosity using ratio of area without presence of nanofibers to total image area as well as nanofiber diameter. Alignment of PVDF-TrFE nanofibers was quantified in ImageJ (version 1.53k) using previously established methods (Eq 2) [127]. Briefly, cropped SEM fiber images (512 x 340 pixels) were subjected to Fast Fourier Transform (FFT) analysis to generate polar plots and respective alignment index. Groups utilized for alignment comparison (n=5) consisted of 3 h electrospun PVDF-TrFE nanofibrous scaffolds under no treatment and annealing at 80°C, 100°C, 120°C, and 140°C. Five different images (n=5) were used for each condition to quantify fiber alignment.

### 5.6 Mechanical Testing & Atomic Force Microscopy

Annealed PVDF-TrFE scaffolds, in groups of four for each annealing temperature of NT 80°C, 100°C, 120°C, and 140°C, were tested for tensile properties using a custom testing machine (100R6; TestResources, Shakopee, MN, USA). These scaffolds, sized at 1.5 cm × 3 cm, were elongated parallel to the fiber orientation at a rate of 0.167 mm/second until failure. The tensile testing captured Young’s modulus and ultimate tensile strength for each scaffold group, utilizing MtestWR software to assess the mechanical integrity of PVDF-TrFE across different applied force.

Annealed PVDF-TrFE scaffolds at multiple temperatures were used for stiffness measurement with an AFM (NanoWizard IV, JPK Instruments). The AFM head with a silicon nitride cantilever (CSC37, k = 0.3-0.8 N/m, f = 20-40 kHZ, MikroMasch) was mounted on a fluorescence stereo microscope (M205 FA, Leica) coupled with a Z-axis piezo stage (JPK CellHesion module, JPK Instruments), which allows the indentation measurement up to the depth of ∼ 100 μm. The sample slide was then placed onto the AFM stage to take AFM micrograph images in non-contact and Qi mode, and force-distance curves in a 10×10 μm square were measured from each nanofiber under ambient conditions for n=100 reads. The Young’s moduli (E, Pa) of the nanofibers were determined by fitting the obtained force-distance curves with the modified Hertz model [128] and resultant stiffness values were analyzed.

### 5.7 Fourier transform infrared spectroscopy and X-ray diffraction

Fourier Transform Infrared Spectroscopy (FTIR) (Nicolet 6700 with Smart Orbit diamond ATR, Thermo Fisher Scientific) and X-ray Diffraction (XRD) (MINIFLEX, Rigaku) were utilized to evaluate and analyze the crystallinity and crystalline structure of annealed PVDF-TrFE scaffolds from four annealed electrospun PVDF-TrFE scaffolds of each annealing temperature group. Attenuated Total Reflectance (ATR) FTIR was employed, covering a range of 4000–400 cm^−1^ with a resolution of 4 cm^−1^. The Lambert-Beer law was applied to approximate the β-phase content (Eq 3). XRD analyses were conducted by irradiating annealed PVDF-TrFE scaffold samples with monochromatic Cu Kα radiation at a scan rate of 0.013°/s, with 2θ ranged between 15° and 25° to capture the peak beta crystallinity relative intensity.

### 5.8 Piezo response Voltage Measurements

PVDF-TrFE scaffolds, including both unannealed scaffolds and those annealed at 140°C, were placed into piezo-responsive chips. This was achieved by incorporating a cantilever beneath the scaffold of size 1.5 x 3 cm. The scaffold was then affixed to a glass slide using a layer of conductive silver paint (Ted Pella Inc.) serving as an adhesive for electrodes on opposite sides of the slide. These opposing electrodes were connected to an x10 voltage amplifier (Charge Amplifier for Piezo Sensor, Polyk Technologies), which in turn was linked to an oscilloscope (Infinium S-Series Oscilloscope, Infinium) for voltage measurement (n=6) in response to mechanical stimulation. The scaffolds underwent deformation approaching ultimate tensile forces through maximal scaffold deformation for measurable electrical signals to be captured using the oscilloscope and were filtered for analysis using MATLAB. Peak to peak electrical responses were then calculated after filtering and were compared between NT and a calculated filtering of noise and the extraction of comparative signals to evaluate peak-peak electrical response.

### 5.9 Cell Culture

For cell culture preparation, scaffolds underwent an additional 30-minute UV light sterilization and were then rinsed with 70% ethanol. This was followed by three PBS washes and an overnight soak in cell culture medium. To maintain the scaffolds submerged during cell culture, polytetrafluoroethylene (PTFE) O-rings (Wilmad Labglass) were positioned on the scaffolds before cell seeding. RT4-D6P2T Schwann cells (ATCC) and NIH 3T3 fibroblasts (ATCC) were cultured in high glucose Dulbecco’s modified eagle medium (DMEM) (SH30022) (GE Healthcare) supplemented with either fetal bovine serum (10%) (Thermo Fisher) for Schwann cells or bovine calf serum (10%) (Thermo Fisher) for fibroblasts and pen/strep (1%) (Thermo Fisher) at 37°C using CO2 (5%) and 95% relative humidity. Schwann cells and fibroblasts were seeded at a density of 100 cells per mm² on NT and 140°C annealed PVDF-TrFE scaffolds sized at 2 cm x 2 cm post sterilization procedure, affixed to 18 mm x 1.5 mm circular coverslips, and placed into 12- well tissue culture plates (Corning, Corning, NY). Cells were grown to sub confluence before passaging via phosphate buffered saline (PBS) (Thermo Fisher) wash and dissociation by trypsin (0.25%) in versine (Gibco) solution.

### 5.10 Cell Viability on Annealed Scaffolds

RT4-D6P2T Schwann cells and NIH3T3 fibroblasts were cultured onNT and 140 °C annealed scaffolds at a seeding density of 50 cells per mm² to evaluate metabolic activity. To quantify this activity, the colorimetric MTT Viability Assay from MilliporeSigma (Calbiochem, Fisher Scientific) was employed, establishing a standard absorbance curve for living cells. This process, which measures the reduction of MTT by cellular metabolic enzymes, enables the assessment of metabolic activity via the optical density detected at 540 nm (OD540nm) with a microplate reader (Bio-Rad). This optical density serves as an indirect indicator of cell viability. The metabolic activity of both cell types on the scaffolds was assayed at three time points: 4, 24, and 48 hours post-seeding, allowing for a comprehensive evaluation of cell viability over time.

### 5.11 In Vitro Immunofluorescence staining and microscopy

Cultured Schwann cells were fixed onto nanofiber scaffolds with 10% formaldehyde at room temperature for 15 minutes. Incubations with both primary and secondary antibodies took place overnight at 4 °C in phosphate buffered saline with 1% bovine serum albumin. Hoechst was used as a nuclear stain. Mouse anti-GFAP (Thermo # MA5- 12023) was used at a dilution of 1:700, and rabbit anti-MKI67 (Thermo # RM-9106) was used at a dilution of 1:500. Scaffolds were mounted to slides for imaging. Images were captured on a Nikon Eclipse Ti and a Nikon AXR Confocal. Analysis was done using the Nikon Elements Imaging Software suite (Nikon).

### 5.12 Western Blot

Protein concentration levels of lysates were analyzed using a BCA protein assay and quantified using a colorimetric 96-well microplate reader (Biorad). A total of 4 μg of protein were extracted from each lysate and reduced with 1:1 dilution of 2-mercaptoethanol (Thermo Scientific) and 2× Laemilli Buffer (Biorad). Lysates were heated for 5 min at 96 °C and resolved on an 8–10% SDS-acrylamide gel (Invitrogen). Following electrophoresis, proteins were transferred onto a PVDF membrane using a Trans-Blot Turbo System (Biorad) and semi-dry transfer buffer. Membranes were blocked with a 2 % BSA in TBST (Tris-Buffered Saline, 0.1% Tween-20) solution for 1 h. N-Cadherin Antibody (rat, 1:2000) (13A9), RhoA Antibody (rat, 1:1000) (1A11-4G10), and FAK Antibody (rat, 1:4000) (OTI4D11) (Nouvous) were diluted in blocking solution. Anti-GAPDH (3:1000) monoclonal antibody (mouse) (Invitrogen) was used as a loading control to ensure equal protein loads across all sample lysates. PVDF membranes were incubated in the primary antibody solution at 4 °C overnight. After incubation, membranes were washed with TBST solution between steps and then incubated with either enhanced chemiluminescence (ECL) Rabbit IgG, HRP-linked secondary antibody (GE Healthcare) or enhanced chemiluminescence (ECL) Mouse IgG, HRP-linked secondary antibody (GE Healthcare) for 1 hour at room temperature. Subsequently, membranes were incubated in ECL Western Blotting Substrate (Pierce, Thermo Fisher) for an additional 5 minutes before imaging using the rapid-auto protocol on a ChemiDoc Imaging system (Bio-Rad). Each blot was verified with two additional replicates.

### 5.13 RNA Isolation, Sequencing

Schwann cells (SCs) seeded on both NT and 140°C annealed PVDF-TrFE scaffolds were harvested after 72 hours. The scaffolds were maintained on ice and subjected to repeated vertexing to ensure complete cell detachment. The cells were then suspended, centrifuged, and resuspended in RNAlater stabilization solution (Thermo Fisher, Waltham, MA). Total RNA was isolated using the mirVana miRNA Isolation kit (Thermo Fisher, Waltham, MA). Directional polyA RNA-seq was carried out by the Genomics, Epigenomics, and Sequencing Core at the University of Cincinnati, following established protocols with recent updates [129]. Initially, the quality of total RNA was assessed using a Bioanalyzer (Agilent, Santa Clara, CA) similarly to previous protocols with QC shown in Figure S3 [129].

### 5.14 Bioinformatics Analysis

RNA-seq reads in FASTQ format were first subjected to quality control to assess the need for trimming of adapter sequences or bad quality segments with the use of Rattus_norvegicus (UCSC rn5) reference genome. The programs used in these steps were FastQC v0.11.7, Trim Galore! v0.4.2 and cutadapt v1.9.1 [130]. The trimmed reads were aligned to the reference rat genome version rn5 with the program STAR v2.6.1e [131]. Aligned reads were stripped of duplicate reads with the program sambamba v0.6.8 [132]. Gene-level expression was assessed by counting features for each gene, as defined in the NCBI’s RefSeq database [133]. Read counting was done with the program featureCounts v1.6.2 from the Rsubread package [134]. Raw counts were normalized as transcripts per million (TPM). Differential gene expressions between groups of samples were assessed with the R package DESeq2 v1.26.0 [135]. Gene list and log2 fold change are used for GSEA analysis using GO pathway datasets. Plots were generated using the ggplot2 package and base graphics in R.

### 5.15 Animals & Ventral Subcutaneous Implantation of Nanofiber Meshes

Murine animals were used in biocompatibility determination. Male C57BL/6J inbred mice and aged between eight and ten weeks were used in all transplantation experiments. Mice were housed in the barrier section of the pathogen-free animal vivarium of Cincinnati Children’s Hospital Medical Center (CCHMC). Handling was performed humanely in accordance with the NIH Guide for the Care and Use of Laboratory Animals. C57BL/6J mice were fed standard irradiated chow. Both food and water were provided ad libitum pre- and post-operatively. All experiments were performed with the prior approval of the Institutional Animal Care and Use Committee of CCHMC (Building and Rebuilding the Human Gut, Protocol No. 2021-0060). Mice were anesthetized with 2% inhaled isoflurane (Butler Schein) and their abdomen shaved and prepped in a sterile fashion with alternating applications of isopropyl alcohol and povidine-iodine. 2 cm skin incisions were made along the midline and the skin was separated from the underlying muscle layer through membrane dissection. A 0.7 cm square of sterile nanofiber scaffold was positioned between the muscle and skin layers before closing the incision with a running stitch. Upon closing, all mice were given a single injection of long acting Buprenex (0.05 mg/kg; Midwest Veterinary Supply) for pain management. Survival of mice was followed out to four weeks at which time the mice were humanely euthanized for tissue collection and analysis. No surgical related mortalities were observed within the study.

### 5.16 Ex Vivo Histological staining and microscopy

Tissues were fixed with 4% paraformaldehyde at 4°C overnight, processed and embedded into paraffin blocks. Sectioning was performed at a thickness of 5 μm for staining. Slides were deparaffinized, rehydrated and stained following standard hematoxylin and eosin staining protocols or using a picrosirius red kit (Polysciences, Inc. # 24901-500). Images were captured on a Nikon Eclipse Ti and subsequent analysis was done using the Nikon Elements Imaging Software suite (Nikon). Histological specimens were evaluated by an independent pathologist blinded to sample group.

### 5.17 Statistical Analysis

At least three separate scaffolds were used for each material characterization and in vitro assays. All spun scaffolds were cut up and annealed so there was paired data from same spin, not from separate spins. One way ANOVA and non-parametric tests were conducted with differences represented by p value; p value < 0.05 was considered statistically significant. *: p value < 0.05, **: p value < 0.01, ***: p value < 0.001, ****: p value < 0.0001, ns: not significant.

## Acknowledgements

The authors would like to acknowledge funding for this project by the Department of Defense Congressionally Directed Medical Research Program (Grant #: DM190692), the National Science Foundation Division of Materials Research (Grant #: DMR-2104639), and the National Institute of General Medicine (Grant #: 1R35GM150860-01). The authors would also like to thank Maulee Sheth for help with sample preparation, Dr. Melodie Fickenscher and Mahnoosh Khosravifar for SEM imaging, Cole Farhman for X-ray Diffraction, and Dr. Jha Rashmi, Manoj Yasaswi Vutukuru, and Yara Izhiman for assistance with electrical measurements, Reinaldo dos Santos for tensile testing assistance, Necati Kaval for assistance with Fourier-transform infrared spectroscopy. Additionally, we would like to acknowledge veterinary services staff at Cincinnati Children’s for their support in completing animal work, the Bioinformatics Core at Cincinnati Children’s, and the University of Cincinnati Genetic Core for assistance in RNA sequencing isolation and analysis.

## Data Availability

The RNA-seq datasets generated and/or analyzed during the current study are available in the GEO repository and will be accessible with the accession number provided by the GEO curators. This data will be accessible upon public release on 2028-07-03. The submission includes a metadata file named “seq_template file maksym krutko – Esfandiari Lab.xlsx” and was submitted under the user ID “ronikade@orcid.” The files have been uploaded to the subfolder “uploads/ronikade@orcid_S7U5U4Ci/Leyla_Schwann.” For additional assistance or inquiries regarding the submission, please contact GEO at. If there are any issues with the submission, the corresponding authors can be contacted at.

