## Supporting Information for "Enhanced Piezoelectric Performance of PVDF-TrFE Nanofibers through Annealing for Tissue Engineering Applications"

**Supplementary Materials**

**Table S1: DSC Derived Crystallinity Fraction**

| Groups | Total Crystallinity Fraction (%) |
| --- | --- |
| NT | 15.43 |
| 80 °C | 16.19 |
| 100 °C | 17.29 |
| 120 °C | 22.30 |
| 140 °C | 29.39 |

**Table S2: Summary of Fiber Diameter and Porosity Values**

| Groups | Fiber Diameter (nm) | Porosity (%) |
| --- | --- | --- |
| NT | 533.9 (±99.60) | 0.7 (±0.3) |
| 80 °C | 615.6 (±175.4) | 5.9 (±0.3) |
| 100 °C | 592.7 (±134.2) | 5.0 (±1.7) |
| 120 °C | 579.9 (±120.9) | 6.9 (±2.8) |
| 140 °C | 517.9 (±109.8) | 16.8 (±4.3) |

**Table S3: Summary of Mechanical Characterization of Annealed Groups**

| Groups | Young’s Modulus  (Tensile Testing) (MPa) | Ultimate Tensile Strength (MPa) | Young’s Modulus (AFM) (kPa) |
| --- | --- | --- | --- |
| NT | 11.2 (±1.9) | 11.2 (±2.5) | 56.1 (±9) |
| 80 °C | 1.14 (±0.1) | 2.47 (±0.2) | 76.4 (±12) |
| 100 °C | 3.13 (±1.4) | 2.70 (±0.2) | 109 (±21) |
| 120 °C | 10.71 (±1.9) | 2.28 (±0.2) | 163 (±29) |
| 140 °C | 71.8 (±13) | 17.8 (±1.9) | 160 (±33) |

**Table S4: Significance Summary of Mechanical Characterization of Annealed Groups where (*p<0.05, **p<.01, ****p<.0001)**

| Groups | Young’s Modulus (Tensile Testing) Significance | Ultimate Tensile Strength Significance | Young’s Modulus (AFM) Significance |
| --- | --- | --- | --- |
| NT x 80 °C | ** | * | **** |
| NT x 100 °C | ** | * | **** |
| NT x 120 °C | No Significance | * | **** |
| NT x 140 °C | * | * | **** |
| 80°C x 100 °C | No Significance | No Significance | **** |
| 80°C x 120 °C | * | No Significance | **** |
| 80°C x 140 °C | ** | ** | **** |
| 100°C x 120 °C | * | No Significance | **** |
| 100°C x 140 °C | * | ** | **** |
| 120°C x 140 °C | * | ** | No Significance |

**Table S5: Summary of Piezoelectric Properties of Annealed Scaffolds**

| Groups | FTIR F(β) crystallinity fraction (%) | Maximum Piezo Voltage Response (mV) |
| --- | --- | --- |
| NT | 78.48 (±0.01) | 9.9 (±1.6) |
| 80 °C | 75.53 (±0.01) | N/A |
| 100 °C | 68.70 (±0.01) | N/A |
| 120 °C | 72.36 (±0.02) | N/A |
| 140 °C | 79.50 (±0.01) | 19 (±3.2) |

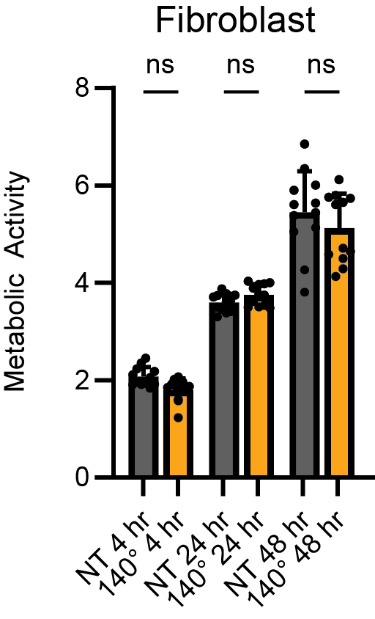

**Figure S1: MTT of Fibroblasts on PVDF-TrFE Scaffolds.** Bar graph of MTT cell metabolism assay comparing NT and 140 group over 48 hours of NIH 3T3 fibroblasts with (n>12 readings).

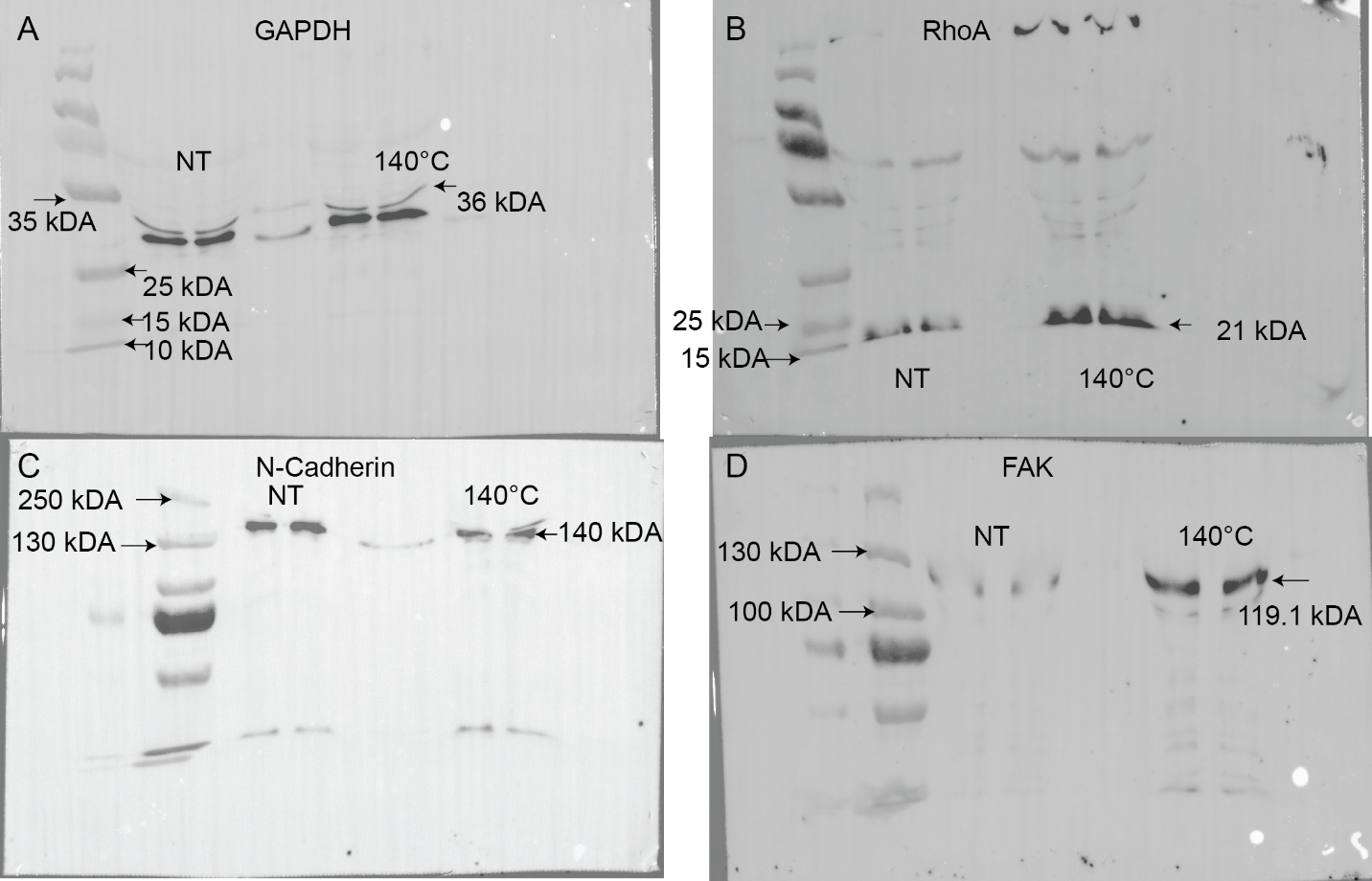

**Figure S2:** Raw Western Blots of Schwann Cell Lysates on PVDF-TrFE Scaffolds

Western blot analysis of Schwann cell lysates after 48 hours on NT and 140 group scaffolds showing the expression levels of A) control protein GAPDH, B) RhoA, C) N-Cadherin, and D) FAK proteins.

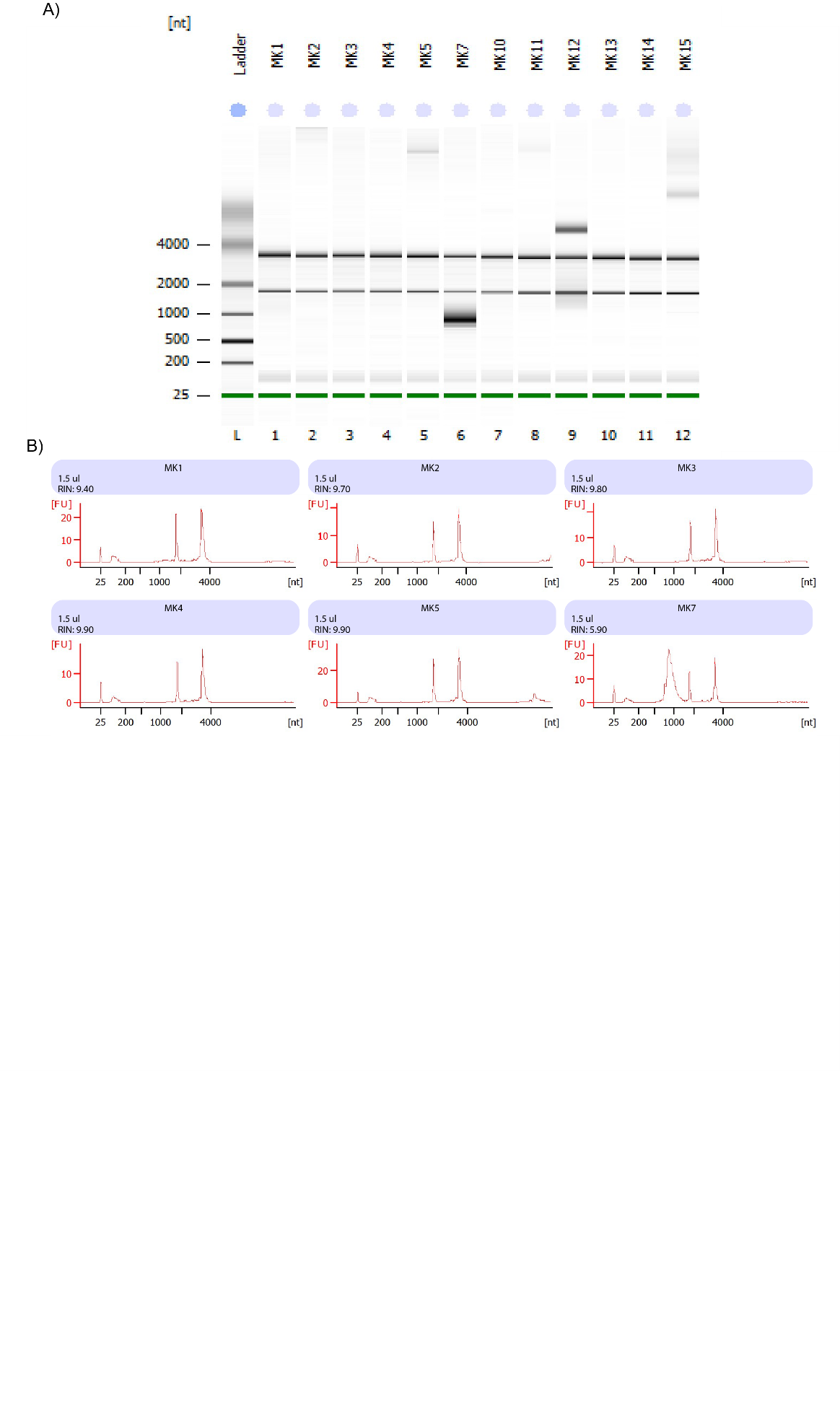

**Figure S3: RNA Quality Control.** A) Electrophoresis run of samples B) Total RNA Analysis ng sensitivity (Eukaryote) with RIN number for samples used in study.
